# Developmental and Environmental Stability of Candidate Reference Genes in the Wild Bee *Ceratina calcarata*

**DOI:** 10.1101/2025.07.28.667322

**Authors:** Zixiao Zhao, Rameshwor Pudasaini, Danielle Kroh, Sandra M. Rehan, Hongmei Li-Byarlay

**Affiliations:** Agricultural Research Development Program, Central State University, Wilberforce, OH, USA; Department of Biology, York University, Toronto, ON, Canada; Department of Agricultural and Life Sciences, Central State University, Wilberforce, OH, USA

**Author notes:** Corresponding author: HL-B. Co-first authors (Zixiao Zhao, Rameshwor Pudasaini). **Author information** ZZ, RP, DK, SMR.

**Keywords:** Small carpenter bee, Gene expression stability, q-PCR, Reference gene, *RPS18*, *RPL8*

## Abstract

Quantitative real-time PCR (RT-qPCR) is a widely used method for measuring gene expression, but its accuracy depends on the use of stable reference genes for data normalization. In this study, we evaluated the expression stability of seven candidate reference genes (*RPS18, RPS5, RPL32, RPL8, EF-1*α, β*-Actin*, and *GAPDH*) in the small carpenter bee *Ceratina calcarata* across developmental stages (larvae, pupae, adults) and different landscape environments (conventional farms, organic farms, and roadside sites). Using four analytical algorithms, GeNorm, NormFinder, BestKeeper, and the comparative ΔCt method, we identified *RPS18* and *RPL8* as the most stable reference genes under varying biological and environmental conditions. These findings were further supported by RefFinder, which integrates results from all algorithms. Our study provides the first validated reference genes for *C. calcarata*, enabling more accurate and reproducible gene expression analysis in this ecologically important wild bee species. This work will support future research in pollinator biology, environmental stress responses, and conservation genomics.

## Introduction

The service provided by pollinators is essential for global ecosystems. About 87.5% of flowering plants worldwide require pollination service by animals [1]. About 75% of the agricultural crops benefit from pollinators, resulting in a 35% increase in yield and generating billions of dollars in value [2,3]. Honey bees (*Apis mellifera*) are the most studied pollinators and have been managed to produce honey and other products. Native wild bees are far more numerous than managed bees. They provide complementary pollination service thought to surpass managed bees, which contribute to the ecological and agricultural stability [4,5]. The small carpenter bees refer to a large group of more than 300 species from genus *Ceratina* (Apidae: Xylocopinae) [6]. *Ceratina calcarata* Robertson is a species native to eastern North America, ranging Florida to the south, Ontario to the north, and Nova Scotia to the east [7]. The species, as well as the sympatric species *C. dupla* Say, can effectively pollinate many fruit, vegetable, and other crops [8,9] and are among the most abundant pollinator species in recent restored land [9,10]. Similar to honey bees, native wild bees also suffered from declining populations, potentially caused by loss of habitat, the use of agrochemicals, land-scape alternation, and parasitism [11,12].

Besides its economic and ecological importance, *C. calcarata* is considered as a subsocial species with features of prolonged maternal care and mother-adult offspring interaction. It also demonstrates traits of facultative sociality such as division of labor and cooperative brood care [13]. The eldest daughters are dwarf in body size and responsible for foraging, guarding and feeding their younger siblings [14–16]. Their roles resemble worker-like behavior in eusocial species and they do not have chance to overwinter or reproduce the next spring. Therefore, *C. calcarata* is an ideal model to study the evolution and mechanism of sociality in hymenopteran insects.

To date, many molecular and genomics approaches have been adopted to study the phylogeny [7,17,18], adaptation [19,20] and reproduction [21] of *Ceratina* species. The reference genome of *C. calcarata* [22] and *C. australensis* [23] has been sequenced and assembled. The transcriptome and metatranscriptome of *C. calcarata* have been sequenced, which revealed the gene and microbiome regulations associated with overwintering [24], maternal and sibling care [15,25,26], social behavior [27], and landscape adaptation [28]. Compared to high-throughput sequencing, real-time quantitative reverse transcription polymerase chain reaction (RT-qPCR) provides a fast and accurate approach to quantify genes [29,30,31]. It involves the reverse transcription of RNA into complementary DNA (cDNA), followed by real-time PCR amplification. This method is cost-effective when a small number of target genes are analyzed from a large number of samples. The calculation of relative expression is relied on internal controls, which are housekeeping genes with constant expression levels across the treatments [32]. However, as a perquisite procedure for studying gene expressions associated with development and adaptation to agricultural landscapes, there is a lack of highly conserved reference genes in *C. calcarata* for RT-qPCRs. Thus, it is a challenge to study target gene expressions in this bee species.

In this study, we tested the expressional stability of seven commonly used reference genes, including four ribosomal binding proteins: ribosomal protein L8 (*RPL8*) and L32 (*RPL32*) binding to large subunits, S5 (*RPS5*) and S18 (*RPS18*) binding to small subunit, a cytoskeleton protein: β-actin (*ACT*), a translation elongation protein: elongation factor 1-alpha F2 (*EF-1α*), and a housekeeping enzyme: glyceraldehyde 3-phosphate dehydrogenase (*GADPH*). We used four methods to compare the stability of each gene: comparative ΔCt analysis [33], NormFinder [34], geNorm [35], and BestKeeper [36]. Further RefFinder was utilized for integrated analysis with incorporating GeNorm, BestKeeper, NormFinder, and ΔCt analysis [37]. Our results present the most stable genes across landscapes and developmental stages, which can be used in studies of gene expression analysis under similar scenarios.

## Materials and methods

### Sample collection and total RNA extraction

*C. calcarata* individuals were collected from multiple sites in Western Ohio during the spring 2024. The sites include three types of landscape: conventional farms with regular applications of pesticides and other agrochemicals, organic areas with limited agrochemical applications, and roadside sites without agricultural activity. Based on their nesting behavior in raspberry (*Rubus idaeus* L.) [14,38], dead raspberry canes with diameters over 7 mm from the previous growing season were cut to 1.2-m sticks and vertically inserted into the soil for about 20 cm in early May to attract *C. calcarata* oviposition (Supplementary figure S2).

After four weeks, the nesting individuals were collected from sticks and visually identified their life stages into larva, pupa and adult stages. The samples were frozen in the dry ice and stored temporarily at −80 °C. Individual total RNA was extracted using TRIzol and purified by ZYMO Direct-zol RNA Miniprep Kit (Zymo Research, Irvine, CA, USA). Genomic DNA contamination was removed by DNase I (Zymo Research) using in-column digestion method. The concentrations of RNA were measured by Qubit RNA BR Assay (Thermo Fisher Scientific, Waltham, MA). The RNA samples were preserved at −80 °C for future use.

### Primer design and RT-qPCR experiments, and efficiency test

The predicted coding sequences of candidate genes were searched from *C. calcarata* genome assembly annotation [22] and confirmed by PCR. The primers were designed using Primer3Plus with qPCR module [39] (Table S1). The first strand cDNA for each sample was synthesized by iScript cDNA Synthesis Supermix (Bio-Rad, Hercules, CA) following the factory protocol. A mixture of oligo(dT) and random hexamers were used to prime the reaction. Samples were normalized to 1 μg total RNA per 20 μl reaction mix. Real time PCR were conducted using PowerUp SYBR Green Mix (Thermo Fisher Scientific) in QuantStudio 3 Real-Time PCR System (Thermo Fisher Scientific) by following procedure: 50 °C for 2 minutes, 95 °C for 2 minutes; 40 cycles of 95 °C for 15 seconds, 60 °C for 30 seconds; then a 60-95 °C melting curve to confirm the specificity of amplification. Three technological replications were incorporated for each sample. To obtain the PCR efficiency of each primer set, qPCR was also performed on 10 X serial dilution of cDNA.

## Data analysis

The cycle threshold (Ct) value of each reaction was obtained by Design & Analysis 2 (DA2) software (version 2.8.0, Thermo Fisher Scientific). Standard curves of Ct were made by liner regression, and the efficiency of each primer set was calculated by formula *Efficiency=10^−1/slope^*. We addressed the expressional stabilities under developmental stages or agricultural landscapes using following algorithms: comparative ΔCt method, NormFinder, geNorm, and BestKeeper. R Package ctrlGene (version 1.0.1) [40] were used to address geNorm and BestKeeper analysis. RefFinder was utilized for integrated analysis with incorporating GeNorm, BestKeeper, NormFinder, and ΔCt analysis [37].

## Results

### Primer specificity and efficiency test

The specificity of primers sets was confirmed by RT-PCR. The results of 1% agarose gel electrophoresis followed by RT-PCR presented the unique bands for each primer set within the expected size range. The single peak in melting curves also confirmed the result. Using cDNA serial dilution as standard, the primers present efficiency from 91.05% to 108.40%, which are within the acceptable range.

### Analysis of candidate reference gene expression

The expression levels of different reference genes of *C. calcarata* under different developmental stages and landscapes is presented in the Fig. 1 a,b. The results reveled a range of Ct values for analysed candidate reference genes varying from 14.83 to 32.01. The Ct values for candidate reference genes varied across developmental stages (larvae vs. adult) and collection sites (conventional vs. organic vs roadside sites). Among all genes, *RPS18* and *RPL8* showed lower Ct variability. However, *GADPH* and *ACT* exhibited greater fluctuation.

**Figure 1.**
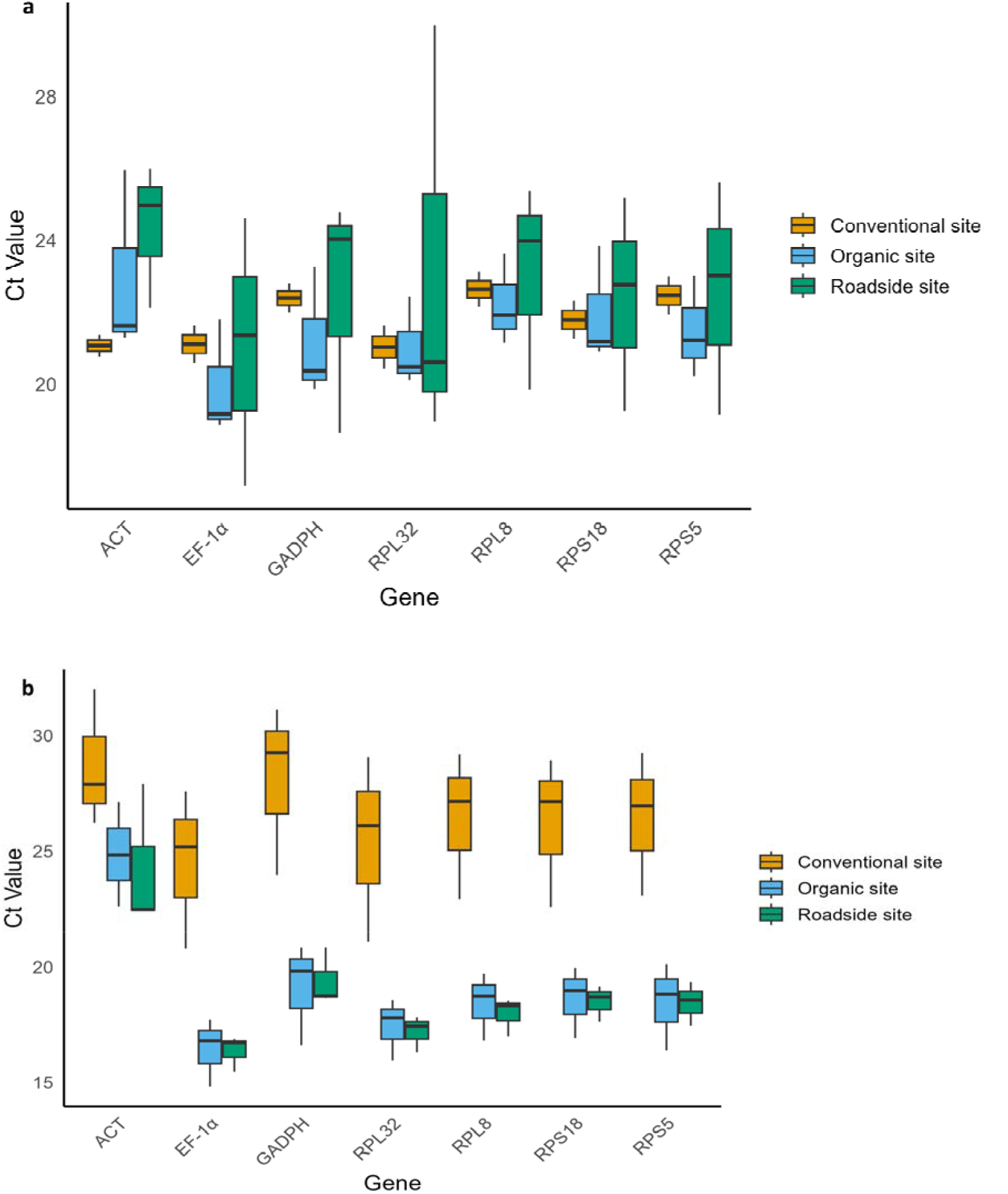
Expression levels of different reference genes of small carpenter bee under different developmental stages and landscapes.

### Stability of candidate reference genes

#### ΔCt method

The ΔCt method utilizes the standard deviations of Ct values to assess gene expression stability of genes. In both larval and adult stages, *RPL8* was the most stable gene among the seven investigated genes. While analyzing the data based on collection sites as organic, roadside and conventional sites, *RPS18, RPS5* and *RPS32* were the more stable genes, respectively (Fig. 2 a-e).

**Figure 2.**
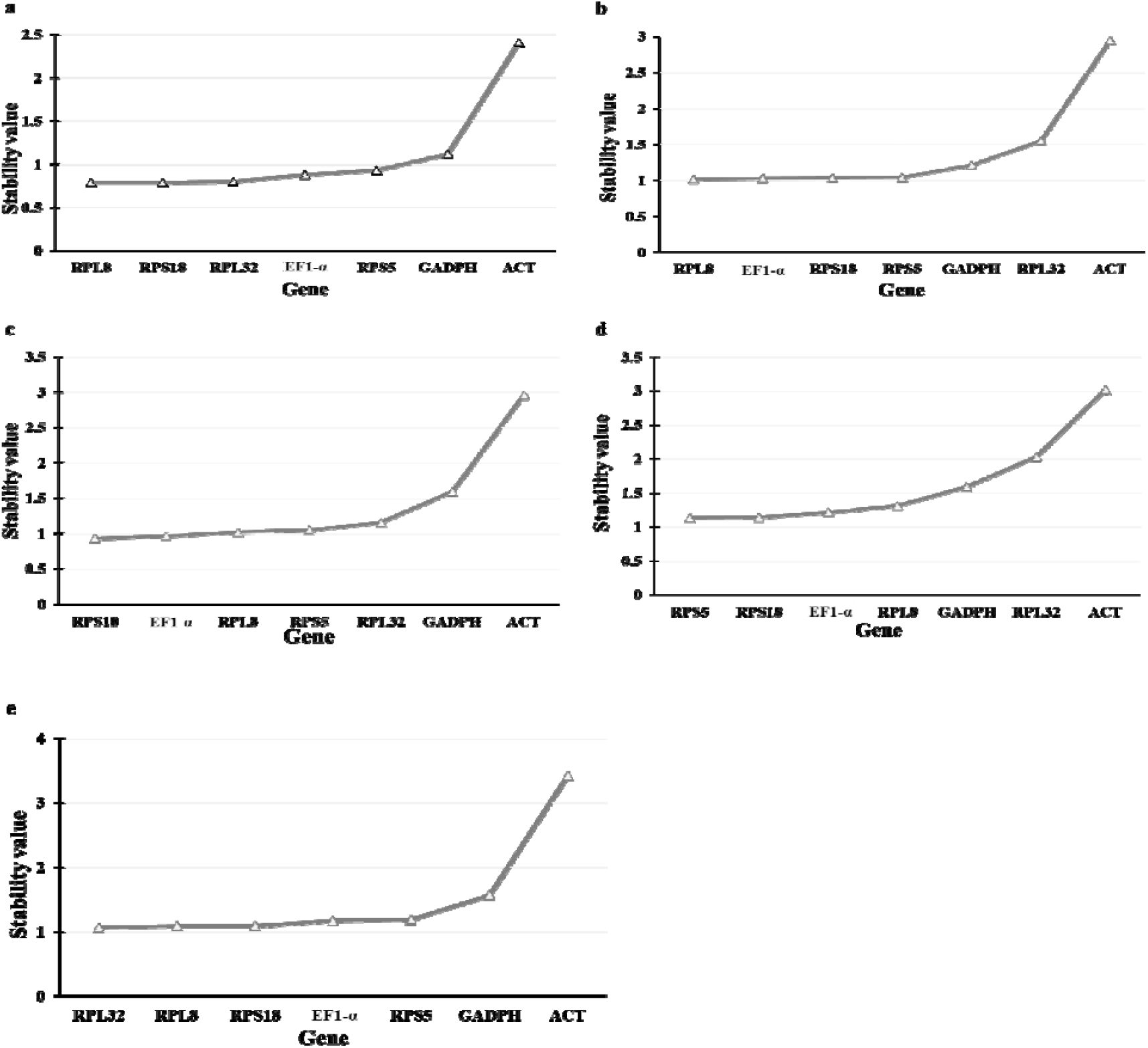
Expression stability of reference genes of small carpenter bee under different developmental stages and landscapes using ΔCt method. (A) Larval stage (B) Adult stage (C) Organic site (D) Roadside site (E) Conventional site

#### GeNorm Analysis

GeNorm analysis used expression stability measurement (M) value based on the average pairwise variation to calculate the stability of expression levels. The M of different candidate reference genes using geNorm are presented in Fig. 3 a-e. *RPL8* followed by *RPL32* were the most stable genes in the larval stage, whereas *RPS18* followed by *RPL8* were the most stable genes in adult stage. However, *RPS18, RPS5* and *RPL8* were the most stable genes based on the organic, roadside and conventional sites of collection, respectively.

**Figure 3.**
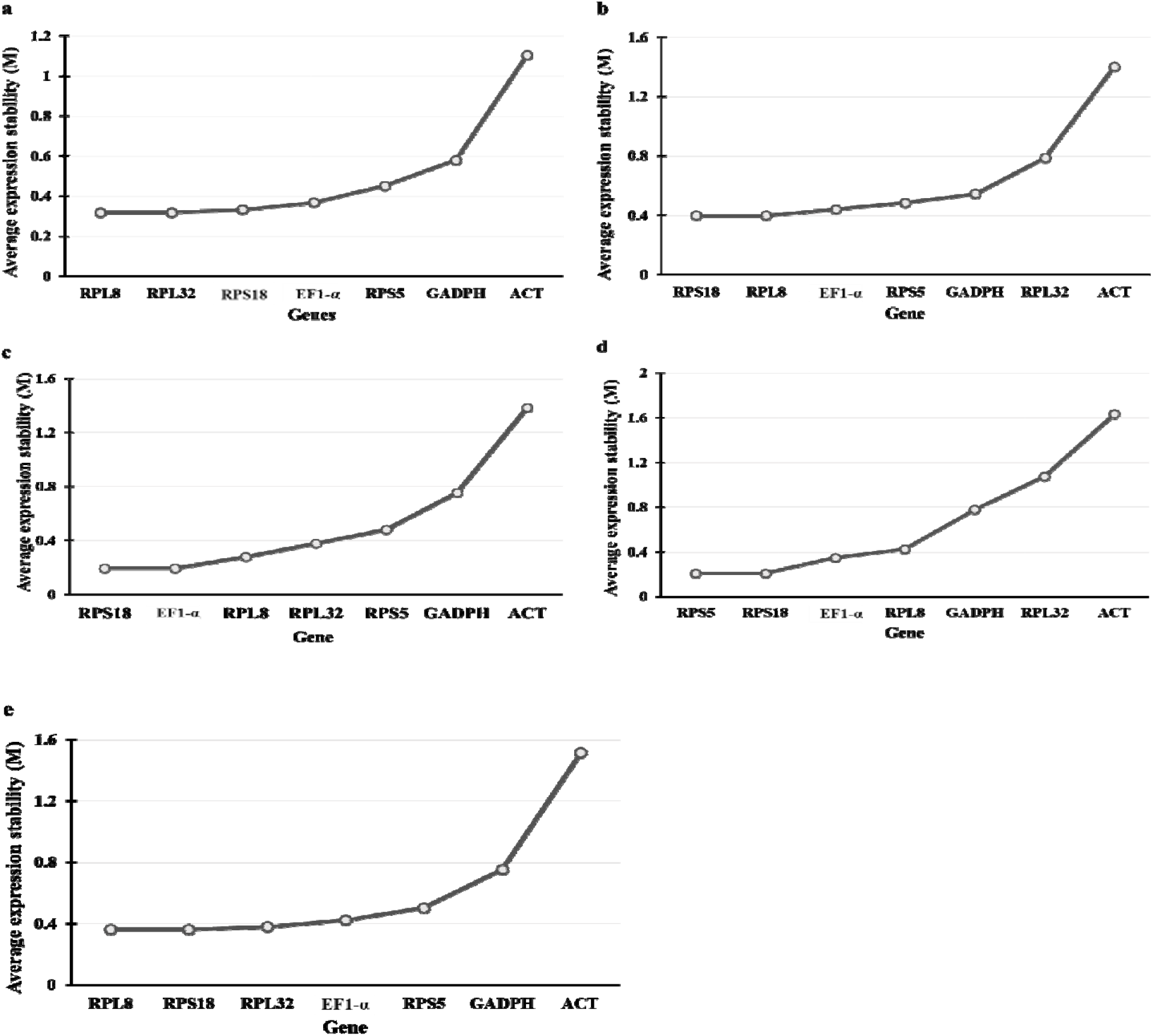
Expression stability measurement (M) of different reference genes of small carpenter bee under different developmental stages and landscapes using geNorm. (A) Larval stage (B) Adult stage (C) Organic site (D) Roadside site (E) Conventional site

#### BestKeeper Analysis

BestKeeper used both the coefficient of variation (CV) and standard deviation (SD) to determine the stability of each candidate reference gene. The stability of a reference gene is considered better if it has a lower CV ± SD value [41]. According to BestKeeper analysis, *ACT* and *RPS18* were the top two most sable genes in the larval stage and RPS18 and RPL8 were the top most stable genes in the adult stage. The most stable candidate reference genes identified were *RPL32* and *RPLS18* on the organic area, *RPS18* and *RPS5* on the roadside area, and *ACT* and *RPS18* on the conventional area (Table 1).

**Table 1.** Expression stability of candidate reference genes in small carpenter bee under different developmental stages and landscapes using BestKeeper analysis.

| Larva |  | Adult |  | Organic site |  | Roadside site |  | Conventional site |  |
| --- | --- | --- | --- | --- | --- | --- | --- | --- | --- |
| Gene | CV ± SD | Gene | CV ± SD | Gene | CV ± SD | Gene | CV ± SD | Gene | CV ± SD |
| *ACT* | 8.53 ± 2.12 | *RPS18* | 8.42 ± 1.66 | *RPL32* | 5.67 ± 1.06 | *RPS18* | 7.69 ± 1.48 | *ACT* | 11.20 ± 2.59 |
| *RPS18* | 10.78 ± 2.02 | *RPL8* | 8.81 ± 1.77 | *RPS18* | 6.35 ± 1.24 | *RPS5* | 8.27 ± 1.59 | *RPS18* | 13.02 ± 2.48 |
| *RP8* | 11.48 ± 2.14 | *RPL32* | 9.41 ± 1.80 | *RPL8* | 6.54 ± 1.28 | *RPL8* | 8.76 ± 1.70 | *RPL8* | 14.09 ± 2.71 |
| *EF-1α* | 11.92 ± 2.01 | *EF-1α* | 10.13 ± 1.83 | *RPS5* | 6.72 ± 1.28 | *GADPH* | 9.14 ± 1.78 | *RPL32* | 15.13 ± 2.72 |
| *RPL32* | 12.23 ± 2.15 | *RPS5* | 10.13 ± 1.96 | *EF-1α* | 6.92 ± 1.21 | *EF-1α* | 9.71 ± 1.69 | *EF-1α* | 15.35 ± 2.68 |
| *RPS5* | 12.69 ± 2.34 | *ACT* | 10.2 ± 2.27 | *GADPH* | 8.26 ± 1.56 | *ACT* | 10.14 ± 2.36 | *RPS5* | 16.52 ± 2.99 |
| *GADPH* | 12.73 ± 2.53 | *GADPH* | 11.31 ± 2.14 | *ACT* | 9.16 ± 2.25 | *RPL32* | 10.82 ± 2.02 | *GADPH* | 16.52 ± 3.23 |

CV: Coefficient of variation, SD: Standard deviation

#### NormFinder Analysis

NormFinder identify the most suitable reference gene by evaluating its expression stability with the lowest value indicating the highest level of stability. The expression stability of candidate reference genes of *C. calcarata* under different developmental stages and landscapes using NormFinder is presented in Fig. 4. *RPS18* and *RPL32* were identified as the most stable candidate reference genes based on the NormFinder analysis.

**Figure 4.**
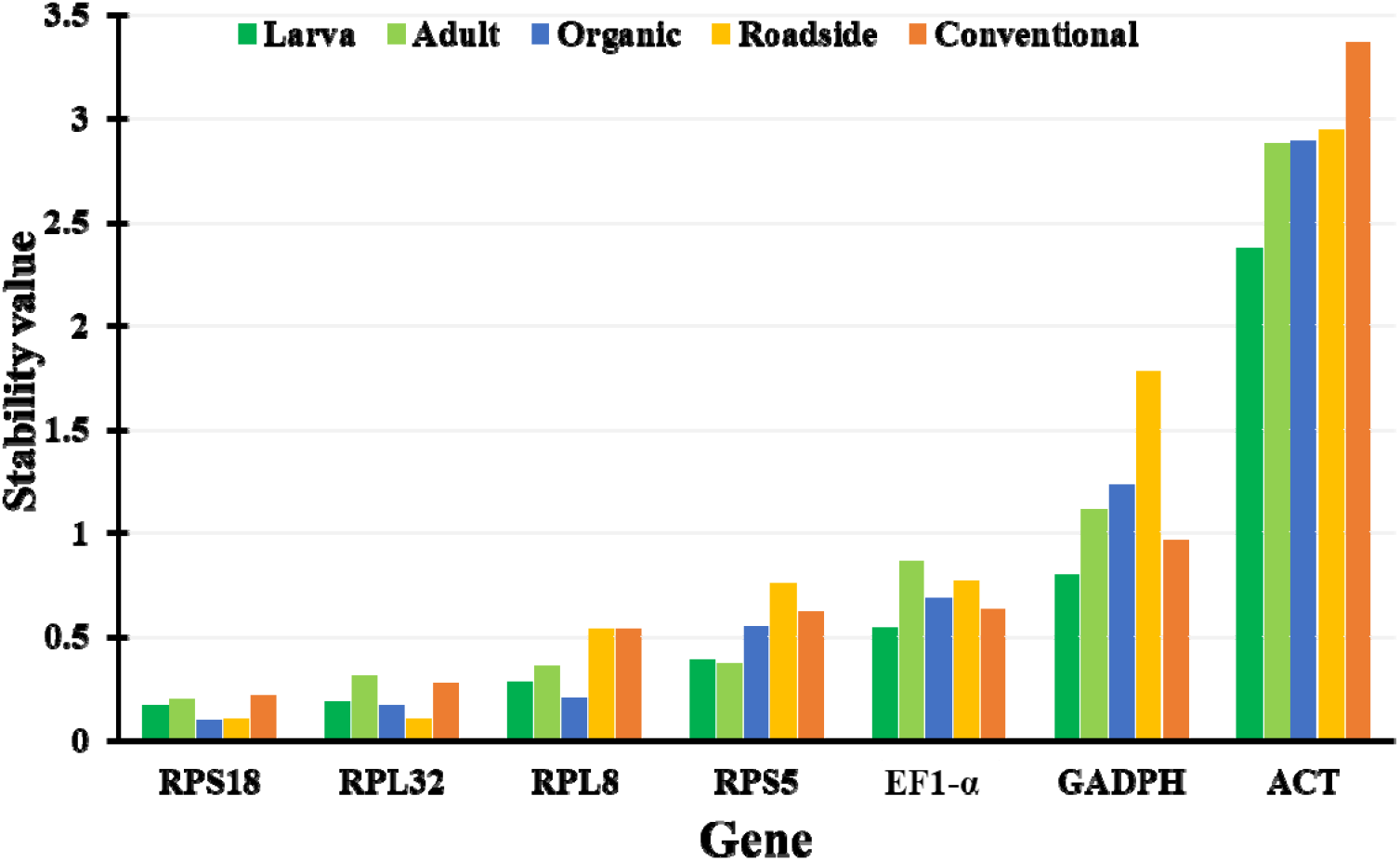
Expression stability of reference genes of small carpenter bee under different developmental stages and landscapes using NormFinder.

#### RefFinder Analysis

RefFinder is an integrated analysis tool used for the validation of reference genes that incorporates several methods including GeNorm, BestKeeper, NormFinder, and ΔCt analysis. Based on the RefFinder, *RPL8* and *RPS18* were the top two most sable genes in the larval and adult stages. *RPS18* consistently ranked among the most stable candidate reference genes across all collection sites, paired with *EF-1*α in the organic site, *RPS5* in the roadside site and *RPL8* in the conventional site (Table 2).

**Table 2.**
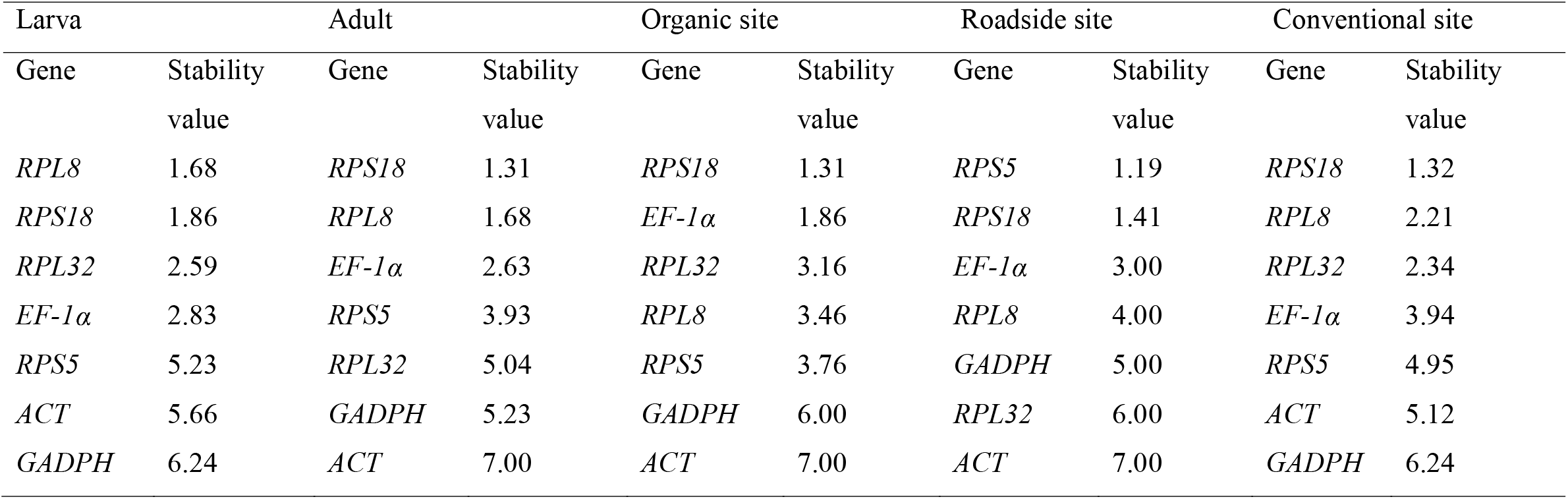
Expression stability of candidate reference genes in small carpenter bee under different developmental stages and landscapes using RefFinder analysis.

## Discussion

*C. calcarata* is considered as an indicator species of healthy ecosystems and important pollinators for natural and agricultural ecosystems [42,43]. This species is commonly used for studying pollinator ecology, behavior, evolution and genomics [14,44]. In this study, we collected larval and adult stages of *C. calcarata* from different habitats, namely organic, roadside and conventional sites. Further we selected seven different candidate reference genes and analyzed their expression stability. To our knowledge, no study has investigated the gene expression stability of *C. calcarata*. Therefore, this study will be an important basis for future studies in *C. calcarata* with implications for native wild bees more broadly.

In this study, we employed widely used analytical tools to evaluate the expression stability of seven candidate reference genes as *ACT, EF-1*α, *GADPH, RPL8, RPL32, RPS5* and *RPS18* in *C. calcarata*. The integrated analysis using RefFinder revealed that *RPS18* consistently ranked among the top two most sable genes across both developmental stages and all habitat types. Similarly, *RPS5* was identified as one of the two most stable genes in larvae, adults and roadside habitat. However, *EF-1*α and *RPL8* were among the other two most stable genes in the organic and conventional site, respectively. In a study of a solitary bee, *Megachile rotundata*, also reported *RPS18*, and *RPL8* as stable reference genes across all life stages and under a variety of environmental conditions [45]. Similarly, a transcriptional study of *C. calcarata* reported significant variation in gene expression associated with overwintering [24]. These findings suggest that *RPS18* and *RPL8* exhibit overall high expression stability and are suitable reference genes for gene expression studies in *C. calcarata* across different developmental stages and habitat conditions.

The expression stability of candidate reference genes varied across developmental stages and habitat types, suggesting that both intrinsic and environmental factors influence gene expression. In other bee species, such as *Euglossa viridissima* also exhibited age-related gene expression patterns [46]. Differences between larvae and adults may reflect distinct physiological processes, such as growth and differentiation in larvae and adults [47]. Additionally, environmental stressors such as pesticide exposure and resource availability may affect gene expression in bees [48]. These findings highlight the need to carefully validate reference genes across both developmental stages and ecological contexts for accurate normalization in RT-qPCR studies.

In conclusion, the present study evaluated the expression stability of seven candidate reference genes in *C. calcarata* across different developmental stages and habitat types. The results demonstrate that gene stability varies with both developmental stages and environmental conditions, underscoring the importance of selecting appropriate reference genes for accurate normalization in RT-qPCR. Our findings provide a valuable resource for future gene expression studies in wild bees and highlight the necessity of validating reference genes under specific conditions.

## Supplementary information

Figure S1. A representative dead raspberry cane used to collect smaller carpenter bees in fields. Table S1. Details of the primers used in this study.

## Acknowledgements

We thank Sakthi Kumaran Subburayalu for their valuable information and technical assistance. We also appreciate Heaven Strachan, Laverne Ambrister, and Keara Clarke for assisting sample collection, and Camiyah Woods for assisting RNA extraction. The research is funded by USDA NIFA awards 2021-38821-34576, 2023-70440-40147, NI231445XXXXG004, and NI241445XXXXG004.

## Data Availability

All raw data and related metadata of this report was deposited in Zenodo, DOI <u>10.5281/zenodo.16414346</u>.

## Author Contributions

**Conceptualization**: Hongmei Li-Byarlay, Zixiao Zhao, Sandra M. Rehan,

**Sample Collection**: Hongmei Li-Byarlay, Zixiao Zhao, Danielle Kroh,

**Formal analysis**: Rameshwor Pudasaini

**Investigation**: Zixiao Zhao, Hongmei Li-Byarlay,

**Resources**: Hongmei Li-Byarlay

**Validation**: Hongmei Li-Byarlay

**Writing – original draft**: Rameshwor Pudasaini, Zixiao Zhao

**Writing – review & editing**: all authors

